# A variant in *SMOC2,* inhibiting BMP signaling by competitively binding to BMPR1B, causes multiple epiphyseal dysplasia

**DOI:** 10.1101/2020.01.27.921825

**Authors:** Feng Long, Lin Li, Hongbiao Shi, Pengyu Li, Shaoqiang Guo, Yuer Ma, Yan Li, Shijun Wei, Fei Gao, Shang Gao, Meitian Wang, Ruonan Duan, Xiaojing Wang, Kun Yang, Ai Liu, Anran Wang, Xiao Chen, Wenjie Sun, Xi Li, Jiangxia Li, Qiji Liu

## Abstract

Previously study showed that SMOC, a matricellular protein, inhibits BMP signaling downstream of its receptor via activation of mitogen-activated protein kinase (MAPK) signaling. In our study, exome sequencing revealed a missense mutation (c.1076T>G, p.Leu359Arg) in EC domain of SMOC2 in a Chinese family with multiple epiphyseal disease (MED). The pathogenicity of this SMOC2 variant was verified by *Smoc2^L359R/L359R^* knock-in mice. Of note, decreasing phosphorylation of SMAD1/5/9 was detected in growth plates and primary chondrocytes from *Smoc2^L359R/L359R^* mice. Furthermore, binding affinity of mutant SMOC2 with collagen IX and HSPG in the extracellular matrix of cartilage were reduced while binding affinity with BMPRIB was intact. In addition, in contrast to previously results, that SMOC2 cannot antagonize BMP activity in the presence of a constitutively activated BMP receptor. These results support that *SMOC2* with p.Leu359Arg variant act as an antagonist of canonical BMP pathway by competitively binding with BMP receptors.

## Introduction

Multiple epiphyseal dysplasia (MED, MIM 132400) is a genotypically and phenotypically heterogeneous skeletal dysplasia and chondrodysplasia that affects epiphysis of long bones^[1, 2]^. MED is characterized by early-onset arthritis, especially in hip and knee joints, which results in joint pain and stiffness, waddling gait in early childhood, and mild to moderate shortness of stature. Eight disease genes, which may be inherited in an autosomal dominant or recessive pattern, have been identified. Autosomal dominant variants include collagen oligomeric matrix protein (*COMP*)^[3, 4]^, collagen type IX α-1 (*COL9A1*)^[5]^, collagen type IX α-2 (*COL9A2*)^[6]^, collagen type IX α-3 (*COL9A3*)^[7]^, matrilin-3 (*MATN3*)^[8]^, and collagen type II α-1 (*COL2A1*) ^[9]^. Autosomal recessive variants are the sulfate transporter gene (*SLC26A2*) and calcium-activated nucleotidase-1 (*CANT1*)^[10, 11]^. All proteins encoded by the known MED-associated genes are involved in maintaining the structural integrity of the cartilage extracellular matrix (ECM). All variants in these genes account for the molecular basis of about 70% of MED cases ^[11]^. However, a number of MED cases have no identifiable genetic mutation, and additional genetic etiologies of MED remain to be identified.

SPARC-related modular calcium binding 2 (SMOC2) and SMOC1, its closest homolog, are members of the protein family BM-40 (also known as secreted protein acidic and rich in cysteines [SPARC] or osteonectin) ^[12, 13]^. BM-40 is a prototypic collagen-binding matricellular protein that participates in regulating cell–matrix interactions, in particular influencing bone mineralization, wound repair and other biological functions. The BM-40 family of modular extracellular proteins is characterized by a follistatin-like (FS) domain as well as an extracellular calcium-binding (EC) domain with two EF-hand calcium-binding motifs. SMOC1 and SMOC2 share a common domain organization, containing one FS domain, one EC domain, two TY domains and one SMOC domain, a novel domain with no known homologs. Increasing studies have shown that both proteins interact with the receptor-mediated signaling of several growth factors and play diverse roles in physiological processes involving matrix assembly and extensive tissue remodelling.

In this study, we investigated a family with autosomal dominant MED; we detected a heterozygous c.1076T>G (p. Leu359Arg) missense mutation in *SMOC2* on exome sequencing. To determine the in vivo pathophysiologic mechanism of SMOC2 mutation, we generated c.1076T>G knock-in mice. In contrast to wild-type mice, mutant mice show a dysplastic tibial growth plate with disordered cells in the proliferative zone and expanded hypertrophic zones. In vivo experiments further demonstrated that decreased heparan sulfate proteoglycan(HSPG)-binding ability of mutant *Smoc2* increased SMOC2 level in ECM, which competed to bind with bone morphogenetic protein receptor (BMPR) and inhibit the BMP–Smad1/5/9 signaling pathway. Co-immunoprecipitation supported that mutant SMOC2 lacked the ability to interact with collagen IX but retained the ability to interact with COMP and MATN3.

## Results

### Pedigree and clinical findings of the family with MED, which caused by a c.1076T>G missense mutation in *SMOC2*

In the four successive generations of the family with MED, 8 patients were affected, as confirmed clinically and radiologically; the male/female ratio was close to 1, which suggested an autosomal dominant inheritance pattern (Figure. 1A). The proband IV-9 was a 4-year-old girl with knee pain and stiffness. Joint pain affecting the knee joint occurred after exercise and she had difficulty rising from the floor. Radiographs showed ossification of the femoral and tibial epiphysis that did not proceed homogeneously from the centre to the periphery (Figure. 1B). All the other patients had a history of pain and stiffness in joints and a shorter stature than average (Table 1).

**Figure 1.**
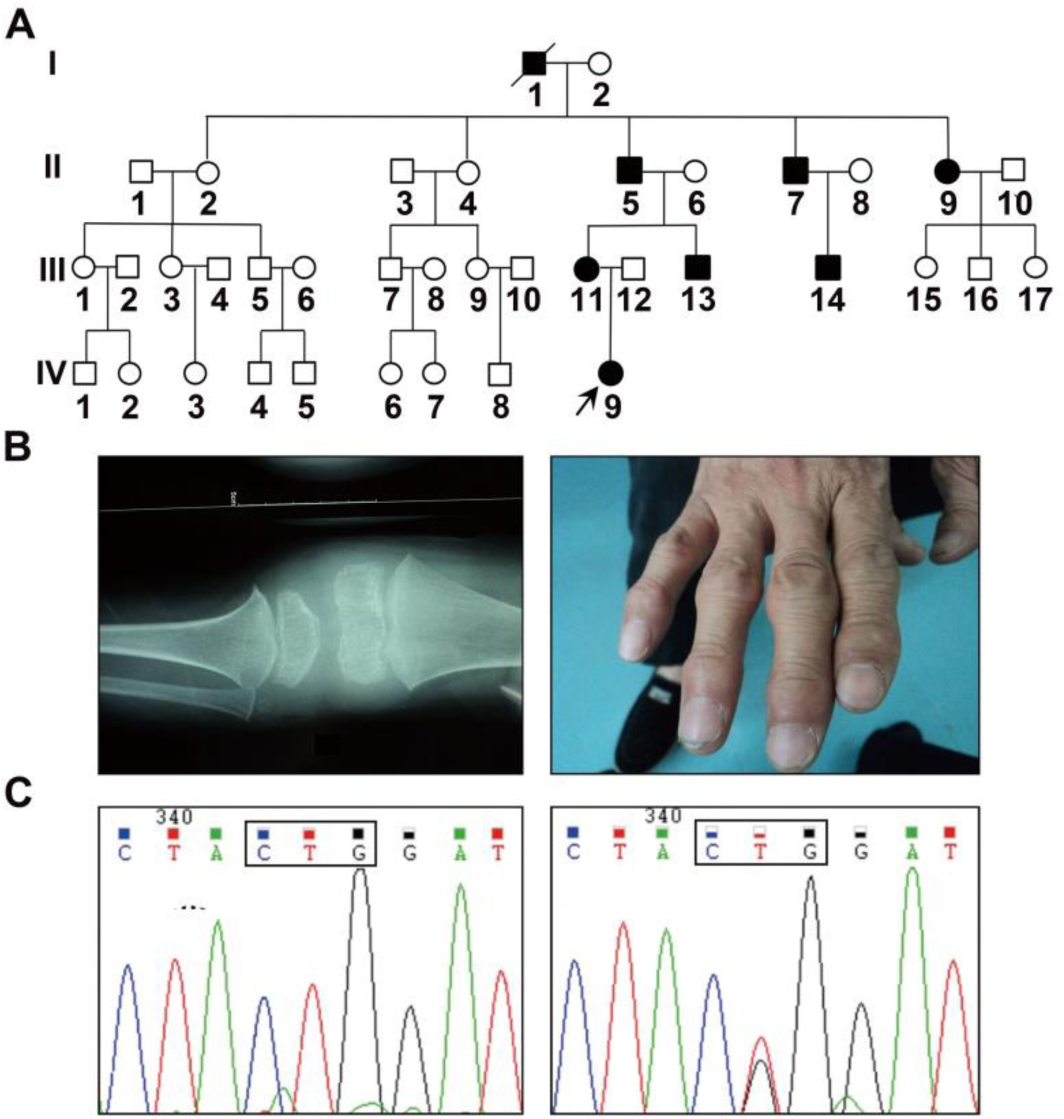
Identification of a mutation in *SMOC2* in a Chinese MED family. A: Pedigree of a Chinese family with autosomal dominant MED in this study. B: Clinical photographs and radiographs of the affected individuals in this family. The picture in the left is anterior-posterior (AP) view of the swelling knee joints of IV9, which showed swelling joint and uneven bone density in epiphysis. The picture in the right is clinical photographs of II7, which showed the swelling finger joints. C: Partial *SMOC2* sequence showed the heterozygous c.1076T>G, p.Leu359Arg mutation in exon 11 of the *SMOC2* gene in the affected family members compared to unaffected family members (normal). Mutant codon (CTG to CGG) is marked by a black box.

**Table 1.** Clinical and Radiological Findings in the Pedigree

| Patient<br>(sex) | Age at Onset<br>(years) | Current Age<br>(years) | Height<br>(cm) | Joint<br>Pain | Clinical Findings |  | Radiological Findings |  |  |  |  |
| --- | --- | --- | --- | --- | --- | --- | --- | --- | --- | --- | --- |
|  |  |  |  |  | Joint<br>Limitation | Other | Epiphyses |  |  |  |  |
|  |  |  |  |  |  |  | Flat | Small | Irregular | OA | Other |
| I1(M) | U | D | 158 | K | K |  |  |  |  |  |  |
| II5(M) | U | 58 | 155 | K, HA | K, HA |  |  |  |  |  |  |
| II7(M) | U | 60 | 160 | K, HA | K, HA |  |  |  |  |  |  |
| II9(F) | U | 53 | 145 | K, H,<br>HA | K, H, HA |  |  |  |  |  |  |
| III11(F) | 5 | 33 | 148 | K, H | K, H |  |  |  |  |  |  |
| III13(M) | 8 | 30 | 158 | K | K |  |  |  |  |  |  |
| III14(M) | 6 | 31 | 159 | K | K |  | K |  |  | K | P<sup>a</sup> |
| IV9(F) | 3 | 9 | 119 | K |  |  | K |  |  | K | K<sup>b</sup> |

Note: U= unknown, D= dead, K= knees, HA= hands, H=hips, P= patella, a = dislocation, b= non-uniform bone mineral density

To find the pathogenic gene, we performed exome sequencing of DNA from patients II9, III14 and IV9 and found 9 variants (Table 2). *PABPC3* was not confirmed by Sanger sequencing in this family, and another 7 variants, except *SMOC2*, could be found in the normal population in the ExAC database (Table 2). Only the NM_001166412.2:c.1076T>G, p.Leu359Arg (L359R) mutation in exon 11 of *SMOC2* could not be found in the normal population. We confirmed a complete cosegregation of the 1076G allele with MED in this family by Sanger sequencing (Figure. 1C). Amino acid sequence alignment of SMOC2 in 12 species showed that the leucine at position 359 was highly conserved (Figure1 - figure supplement 1).

**Table 2.** Selected Candidate Genes and the Allele Frequency in Normal Population

| Gene | Mutation Site | Allele Frequency |
| --- | --- | --- |
| *INSL5* | Exon2: c.197A>G; p. Q66R | 0.00003305 |
| *MRPS5* | Exon5: c.407G>A; p. R136H | 0.0001077 |
| *ANXA5* | Exon4: c.148C>T; p. R50C | 0.00004944 |
| *KAT6B* | Exon11: c.1708C>T; p. R570C | 0.00001648 |
| *MYO15A* | Exon29: c.6316G>A; p. E2106K | 0.000008500 |
| *LRRC48* | Exon4: c.170G>A; p. R57H | 0.00003321 |
| *DSCAM* | Exon32: c.5504C>T; p. T1835M | 0.000008279 |
| *PABPC3* | Exon1: c. 975-979del;p.V325fs | NA |
| *SMOC2* | Exon11: c.1076T>G; p. L359R | NA |

### *Smoc2^L359R/+^* and *Smoc2^L359R/L359R^* mice developed short-limbed dwarfism

In order to study the in vivo pathophysiologic mechanism of *SMOC2* mutation, we generated c.1076T>G knock-in mice (Figure 2 - figure supplement 1A). However, normal Mendelian ratios were not observed in the offspring of all matings. Rare homozygous mutant mice were generated (Table 3). The genotypes of the offspring were confirmed by Sanger sequencing (Figure 2 - figure supplement 1B). The wild-type mice were labeled *Smoc2^+/+^*, heterozygous mutant *Smoc2* mice *Smoc2^L359R/+^,* and homozygous mutant *Smoc2* mice *Smoc2^L359R/L359R^*.

**Table 3.** The number of offspring in different genotypes produced by different mating patterns

| Mating |  | Offspring |  |  |
| --- | --- | --- | --- | --- |
| Male | Female | *Smoc2*<sup>+/+</sup> | *Smoc2*<sup>L359R/+</sup> | *Smoc2*<sup>L359R/L359R</sup> |
| *Smoc2*<sup>+/+</sup> | *Smoc2*<sup>L359R/+</sup> | 11 | 9 | 0 |
| *Smoc2*<sup>L359R/+</sup> | *Smoc2*<sup>+/+</sup> | 28 | 22 | 0 |
| *Smoc2*<sup>L359R/+</sup> | *Smoc2*<sup>L359R/+</sup> | 48 | 111 | 12 |
| *Smoc2*<sup>L359R/L359R</sup> | *Smoc2*<sup>L359R/L359R</sup> | 0 | 0 | 0 |

The gross skeletons of *Smoc2^L359R/+^* and *Smoc2^L359R/L359R^* mice were normal as compared with *Smoc2^+/+^* mice at birth. At postnatal day 30 (P30), obvious differences were observed. The body lengths of *Smoc2^L359R/+^* and *Smoc2^L359R/L359R^* mice were 7.2% and 6.5% reduced as compared with *Smoc2^+/+^* mice; the weights of *Smoc2^L359R/+^* and *Smoc2^L359R/L359R^* mice were 2.2% and 9.7% reduced as compared with *Smoc2^+/+^* mice (Figure. 2A, C). At P63, the difference in body length among all the mice did not expand. However, the weights of *Smoc2^L359R/+^* and *Smoc2^L359R/L359R^* mice were 10.8% and 12.1% reduced as compared with *Smoc2^+/+^* mice (Figure 2 - figure supplement 2A). At P30, the femur and tibia length of *Smoc2^L359R/+^* mice was approximately 7% and 5.7% reduced as compared with *Smoc2^+/+^* mice. The lengths of femur and tibia of *Smoc2^L359R/L359R^* mice were more than 17% and 15.5% shorter than those of *Smoc2^+/+^* mice (Figure. 2A, D). At P63, the tibia lengths of *Smoc2^L359R/+^* and *Smoc2^L359R/L359R^* mice were 7% and 8.5% shorter than those of *Smoc2^+/+^* mice and the reduction in femur lengths was reduced to approximately 5% and 8.1% (Figure 2 - figure supplement 2B).

**Figure 2.**
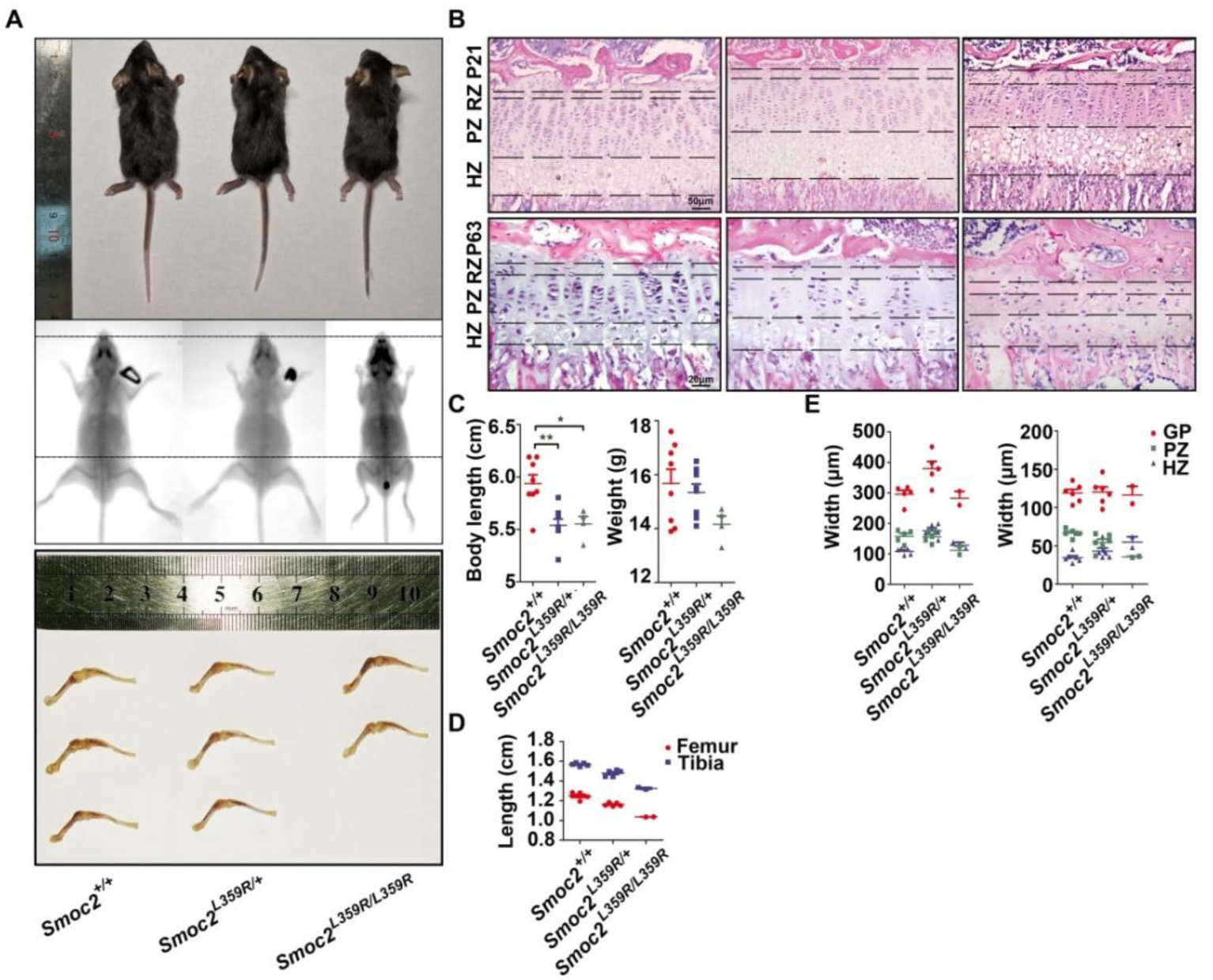
The development of mutant mice was inhibited. A: Photographs and radiographs of *Smoc2^+/+^*, *Smoc2^L359R/+^* and *Smoc2^L359R/L359R^* mice and femurs and tibias of them at P30. Black dotted lines are aligned at the tip of the nose and the top of the pelvis of the *Smoc2^+/+^* mouse. C: Body lengths and weights of all mice were measured at P30. (n=8:8:2, ★ P<0.05, ★★ P<0.01 by t-test.). D: Lengths of femurs and tibias of all mice at P30. (n=5:5:2). B: Hematoxylin and eosin (H&E) staining of tibial growth plates and the hypertrophic zone of tibial growth plate from P21 (the first row) and P63 (the second row) mice. The left column is tibial growth plates of *Smoc2^+/+^*, the middle column is tibial growth plates of *Smoc2^L359R/+^* and the right column is tibial growth plates of *Smoc2^L359R/L359R^*. RZ, resting zone; PZ, proliferative zone; HZ, hypertrophic zone; GP, growth plate. E: Widths of growth plates, proliferative zones and hypertrophic zones of all mice at P21 (left) and P63 (right) (n=5:5:2).

### *Smoc2^L359R/+^* and *Smoc2^L359R/L359R^* mice had disorganized growth plate and abnormal chondrocytes

To determine the effect of mutant *Smoc2* on morphology of tibial growth plates, we used haematoxylin and eosin (H&E) staining of tibial growth plates of all mice at P21 and P63. All tibial growth plates of studied mice were clearly divided into resting, proliferative and hypertrophic zones. At P21, the widths of the growth plates from *Smoc2^L359R/+^* mice became wider because the hypertrophic zones were almost 30% longer than those of *Smoc2^+/+^* mice, but the resting and proliferative zones were still similar to those of *Smoc2^+/+^* mice (Figure. 2B, E). However, at P63, the widths of the hypertrophic zones and ratios of width of hypertrophic zone to that of growth plates tended to be normal (Figure. 2B, E). At P21, as compared with *Smoc2^+/+^* mice, for *Smoc2^L359R/L359R^* mice, proliferative zones were almost 29% shorter and hypertrophic zones were almost 18% wider (Figure. 2B, E). At P63, the proportions increased to 46% and 67% (Figure. 2B, E).

In the tibial growth plates of *Smoc2^+/+^* mice at P21 and P63, the chondrocytes in the resting, proliferative and hypertrophic zones were closely aligned, well organized and arranged along the long axis of the tibia (Figure. 3A). The tibial growth plates of *Smoc2^L359R/+^* and *Smoc2^L359R/L359R^* mice developed progressively dysplastic growth plates from birth. The number of chondrocytes in the tibial growth plates were reduced in these mice than *Smoc2^+/+^* mice (Figure. 3B-D) and they were disorganized and failed to arranged into a column, so hypocellular areas could be observed in the proliferative zones and hyaline cartilage of proximal tibia (Figure. 3A).

**Figure 3.**
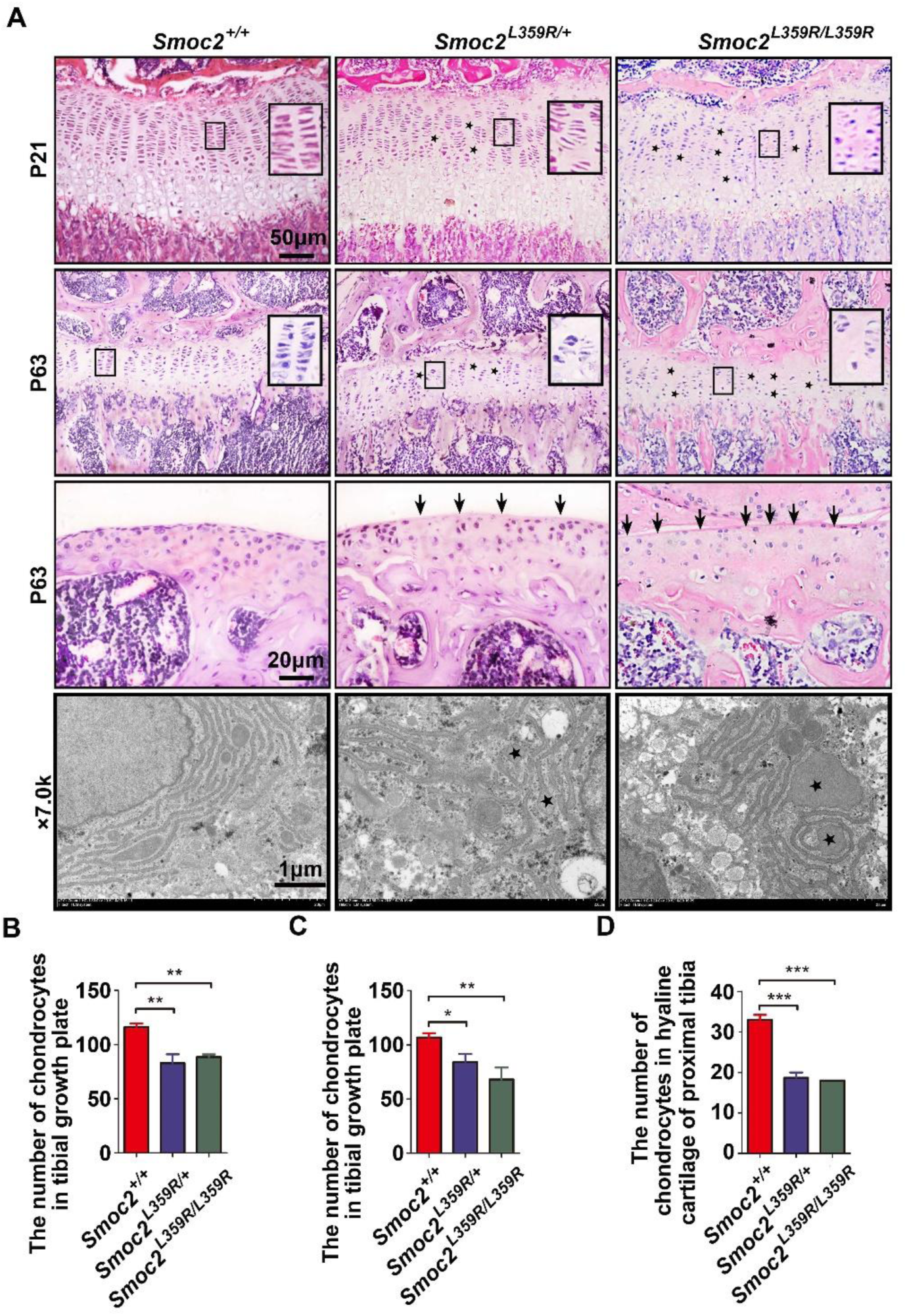
Histological analysis of the tibial growth plate showing growth plate abnormalities. A: The first row is H&E staining of the tibial growth plate of P21 mice showing a disrupted proliferative zone with disorganized columns (black box) and some areas of hypocellularity (black pentagram) in the tibial growth plate from *Smoc2^L359R/+^* and *Smoc2^L359R/L359R^* mice. The second and third rows are H&E staining showing a significantly decreasing cell density (black pentagram) and disorganized columns (black box) in the growth plate and hyaline cartilage (arrowheads) of proximal tibia from *Smoc2^L359R/+^* and *Smoc2^L359R/L359R^* mice compared to *Smoc2^+/+^* mice at P63. The fourth row is ultrastructural analysis of the proximal tibial growth plates of P21 mice by transmission electron microscopy (TEM). Deformed cell nuclear, saw-tooth nuclear membrane and dilated cisternae of rough endoplasmic reticulum in chondrocytes from proximal tibial growth plates of P21 *Smoc2^L359R/+^* and *Smoc2^L359R/L359R^* mice. B-D: Number of chondrocytes in tibial growth plates at P21 and P63 and number of chondrocytes in hyaline cartilage of proximal tibia at P63 (n=5:5:2, ★P<0.05, ★★P<0.01, ★★★P<0.001 by t-test).

On transmission electron microscopy of tibial growth plates of P21 mice, the cisternae of rER in the chondrocytes were well organized and folded. However, in mutant mice, the cell nuclei in chondrocytes were deformed and the nuclear membrane was saw-tooth–shaped. Mutant mice showed some seriously dilated cisternae of rER containing retained proteins in the chondrocytes. Almost all chondrocytes of mutant mice contained seriously dilated cisternae of rER (Figure. 3A).

### *Smoc2^L359R/+^* and *Smoc2^L359R/L359R^* mice showed chondrocyte apoptosis and hypertrophy spatially dysregulated

To determine the mechanism of the expansion of hypertrophic zones in mutant mice tibial growth plates, we determined the relative levels of proliferation, apoptosis and hypertrophy at P21. By using immunohistochemistry with proliferating cell nuclear antigen (PCNA), a marker of proliferation, we found no significant differences in ratio of PCNA-positive cells in both proliferative and hypertrophic zones from the tibial growth plates (Figure. 4A, B). We performed terminal deoxynucleotidyltransferase deoxyuridine triphosphate nick-end labeling (TUNEL) assay on the tibial growth plates of P21 littermates to determine whether decreased apoptosis contributed to the expanded hypertrophic zones in *Smoc2^L359R/+^* and *Smoc2^L359R/L359R^* mice. As expected, mutant mice showed less TUNEL-positive chondrocytes in the hypertrophic zone as compared with *Smoc2^+/+^* mice (Figure. 4A, C). On western blot analysis at P21, Bcl-2 expression was increased and Bax expression was reduced in chondrocytes of mutant mice, which showed the lower level of apoptosis in the mutant mice (Figure. 4D). To test chondrocyte hypertrophy in the growth plates, we used immunohistochemistry and real-time PCR (qPCR) analysis of Ihh. The level of mRNA expression of Ihh was increased in *Smoc2^L359R/+^* mice (Figure. 4E). On immunohistochemistry of cell hypertrophy at P21 (Figure. 4A), the expression area of Ihh (area between the black lines) was wider in *Smoc2^L359R/+^* and *Smoc2^L359R/L359R^* mice than *Smoc2^+/+^* mice. The dysregulated apoptosis and hypertrophy might lead to the expanded hypertrophic zones in the tibial growth plates from mutant mice.

**Figure 4.**
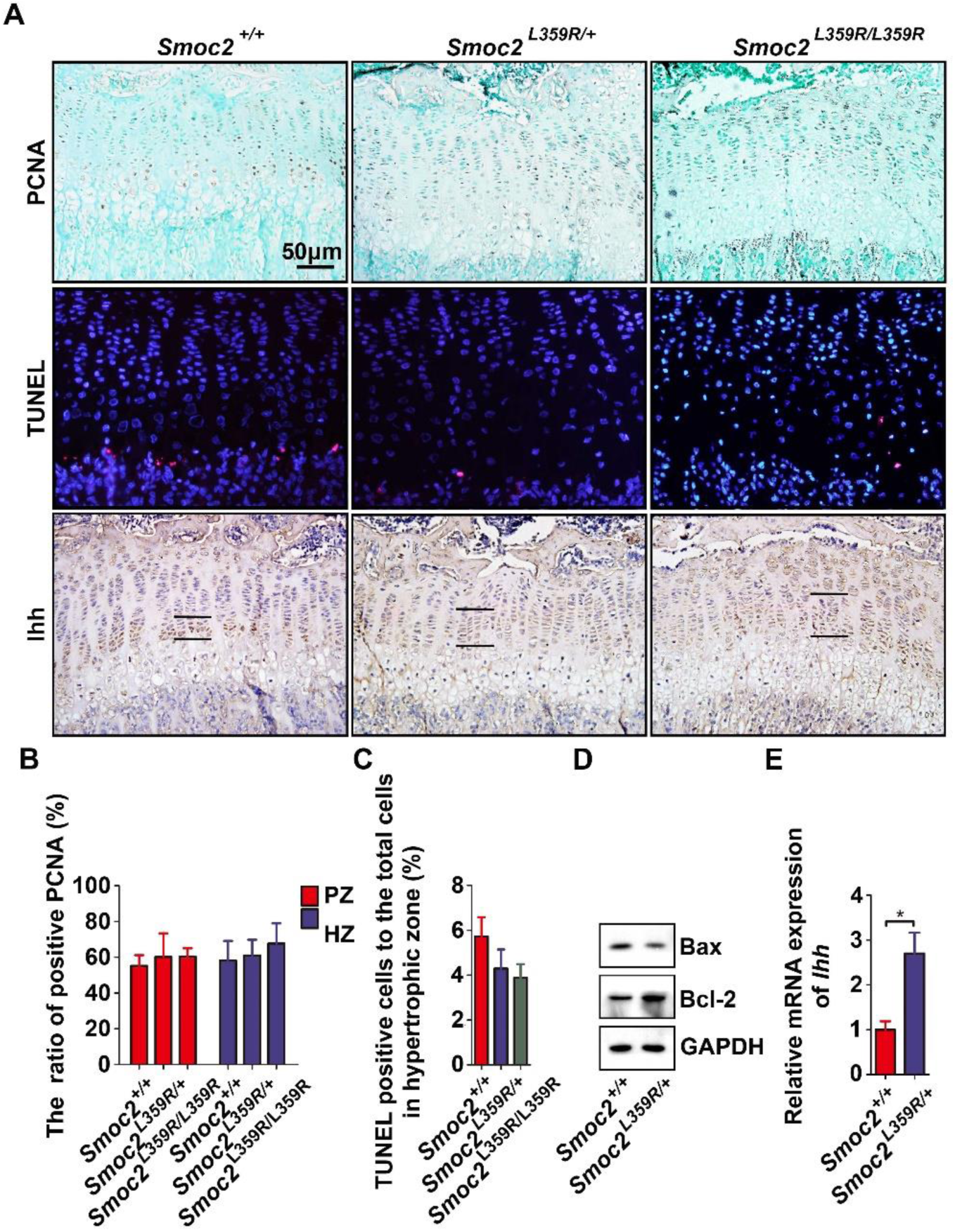
Analysis of proliferation, apoptosis and hypertrophy of chondrocytes in tibial growth plate. A: The first row: Immunohistochemistry of cell proliferation by anti-PCNA antibody on P21 mice. B: The relative proliferation was calculated by comparing the number of PCNA-positive chondrocytes to total number of chondrocytes in the proliferative or hypertrophic zone (n=5:5:2). The second row: Apoptosis was measured in the tibia of P21 mice by TUNEL assay and anti-Bax and anti-Bcl-2 antibodies (D). The relative apoptosis was calculated by comparing the number of apoptotic chondrocytes to total number of chondrocytes in the hypertrophic zone. Nuclei were stained with DAPI (blue) and apoptotic cells were labeled by Cy3 (red). C: Apoptosis of chondrocytes in the hypertrophic zone (n=5:5:2). D: The expression of Bax and Bcl-2 in *Smoc2^L359R/+^* and *Smoc2^+/+^* mice analyzed by western blot analysis. The third row: Immunohistochemistry of cell hypertrophy with anti-Ihh antibodies in P21 mice. The expression area of Ihh (the area between the black lines) in mice. Real-time PCR of the cartilage from proximal tibia of mRNA expression of Ihh (★P<0.05 by t-test) (E).

### Overexpressed SMOC2 and mutant SMOC2 blocked phosphorylation of SMAD1/5/9

Because BMP plays important role in initiation of chondrogenesis and previous study showed that SMOC without EC domain inhibited BMP signalling^[14]^, we analyzed whether mutant SMOC2 inhibited BMP signaling. We found that the number of p-Smad1/5/9 positive cells decreased in primary chondrocytes and tibial growth plates from mutant mice (Figure. 5A, C). The protein level of p-Smad1/5/9 also decreased in mutant mice (Figure. 5B). On western blot analysis of p-SMAD1/5/9 in stably transfected HEK293T cells showed that overexpressed SMOC2 and mutant SMOC2 able to block phosphorylation of SMAD1/5/9 (Figure. 5D) and inhibit the activity of BMP2 (Figure 5 - figure supplement 1A), which signals through SMAD1/5/9.

**Figure 5.**
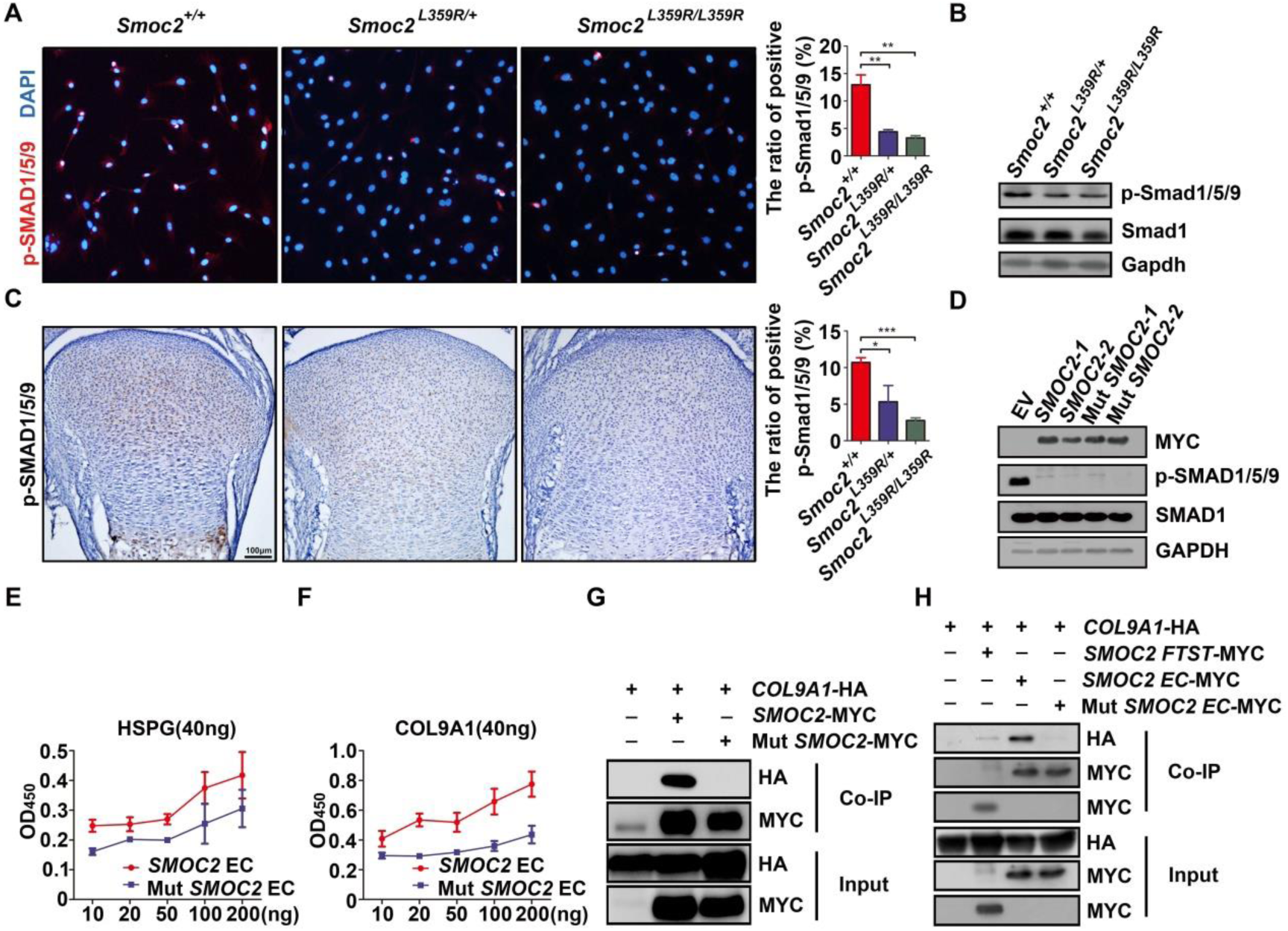
BMP signalling could be inhibited by overexpressed SMOC2 and mutant SMOC2. A: IF analysis of primary chondrocytes from knees showed more positive signals, shown by red fluorescence, in *Smoc2^+/+^* mice compared to *Smoc2L^359R/+^* and *Smoc2^L359R/L359R^* mice. Ratio of positive p-Smad1/5/9 was measured (★P<0.05, ★★★P<0.001 by t-test). B: Western blot analysis of p-SMAD1/5/9 in primary chondrocytes from knees of mice. C: IHC analysis of expression pattern of p-Smad1/5/9 in proximal tibial growth plates from P0 mice. Ratio of positive p-Smad1/5/9 was measured (★★P<0.01 by t-test). D: Western blot analysis of p-SMAD1/5/9 in HEK293T cells stably transfected with wild-type SMOC2 or mutant SMOC2. E-F: Solid phase binding assay to detect binding of SMOC2 EC domain or mutant SMOC2 EC domain to COL9A1 or HSPG. Increasing concentrations of SMOC2 EC domain or mutant SMOC2 EC domain were added to COL9A1-coated or HSPG-coated plates. G-H: Co-IP of HA-tagged COL9A1 by MYC-tagged SMOC2 mutant SMOC2, SMOC2 FTST domain, SMOC2 EC domain and mutant SMOC2 EC domain. HA-tagged COL9A1 could only be detected in the precipitated immunoprecipitation by MYC-tagged SMOC2 or SMOC2 EC domain.

### Overexpressed SMOC2 and mutant SMOC2 inhibited BMP signaling by competitive binding of BMPR1B

In previous study, SMOC1 could inhibit BMP signaling by activating the mitogen-activated protein kinase (MAPK) pathway^[14]^. We used MG132 and BMP2 treated HEK293T cells stably transfected with or without wild *SMOC2* cells (Figure 5 - figure supplement 1B). MYC but not p-SMAD1/5/9 level was rescued and p-SMAD1 (Ser206) level showed no obvious change in cells stably transfected with wild-type *SMOC2-*MYC incubated in MG132 at the indicated doses.

Thomas et al. found that SMOC-EC domain could expand the range of BMP signalling by competitive binding of HSPG with a similar affinity to BMP2^[14]^. We performed solid phase binding assay (SPB) and detected dose-dependent binding of the SMOC2 EC domain or mutant SMOC2 EC domain to HSPG, but the mutant SMOC2 EC domain showed weak binding to HSPG (Figure. 5E). Considering that MED proteins are combined in the ECM, we examined the ability of wild-type and mutant SMOC2 to combine with COMP, MATN3 or COL9A1 by Co-IP and SPB. Mutation on SMOC2 EC domain decreased the ability of binding to COL9A1 (Figure. 5F-H) rather than COMP and MATN3 (Figure 5 - figure supplement 2).

Because SMOC2 did not inhibit BMP signaling by activating the MAPK/ERK pathway, we investigated whether SMOC2 acted by directly binding to the ligand. Western blot analysis showed that inhibition of p-SMAD1/5/9 by SMOC2 could be rescued by activated BMP receptors (Figure. 6A). To clarify the exact BMP receptor binding to SMOC2, we used Co-IP to study the ability of SMOC2 to interact with ACVR1, BMPR1A and BMPR1B. Only BMPR1B could be detected in the precipitate by SMOC2 (Figure. 6B-D). BMPR1B could also be detected in the precipitate by mutant SMOC2 (Figure. 6E). Therefore, both wild-type and mutant SMOC2 inhibited BMP signaling by binding to BMPR1B.

**Figure 6.**
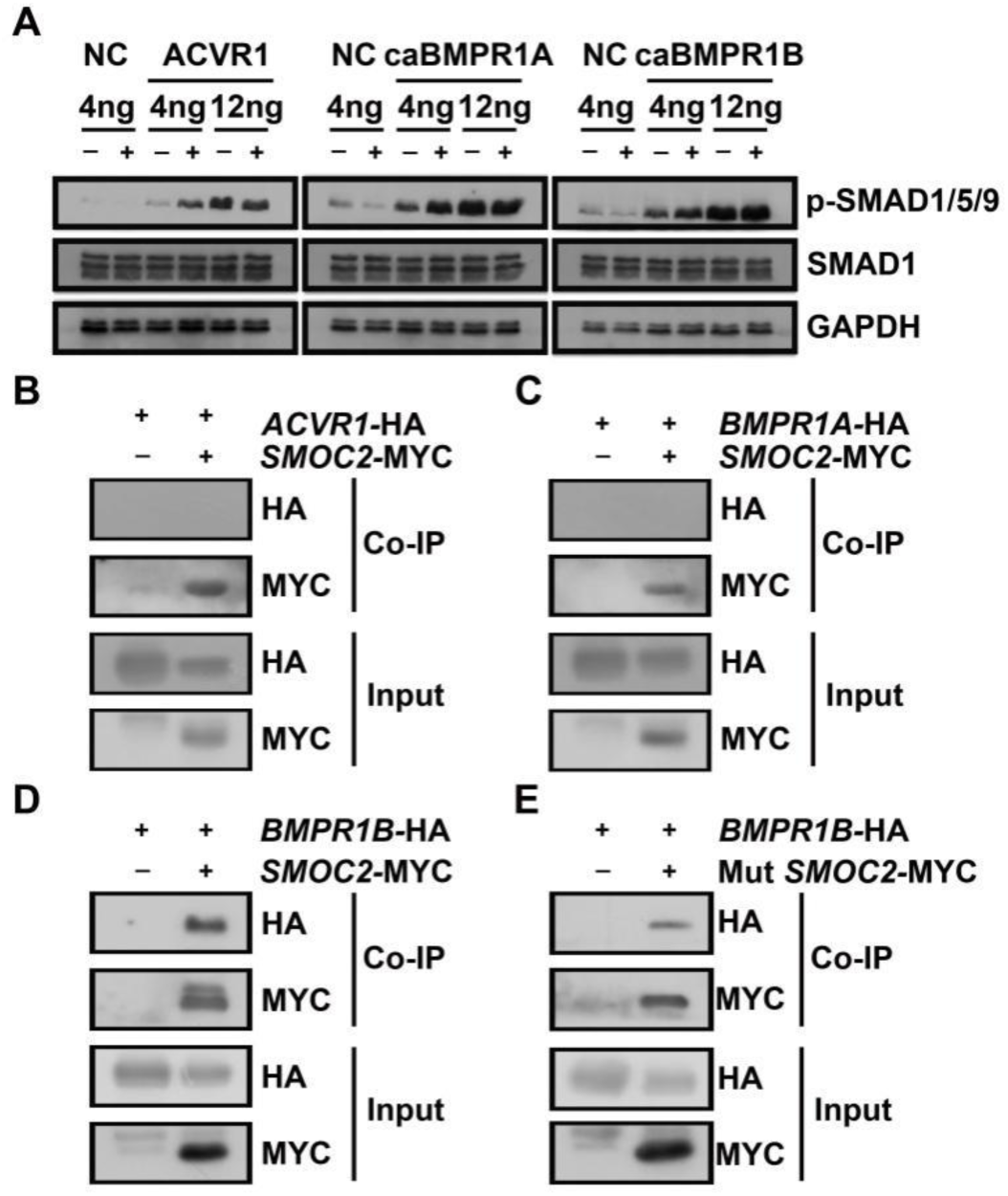
SMOC2 and mutant SMOC2 could bind to BMPR1B. A: Western blot analysis of p-SMAD1/5/9, total SMAD1 and GAPDH in whole cell protein lysates of HEK293T cells transfected with pLVX-IRES-Puro or pLVX-SMOC2 and transiently transfected with pCMV3, pCMV3-ACVR1, pCMV3-caBMPR1A or pCMV3-caBMPR1B. B-E: Co-IP of HA-tagged ACVR1 and BMPR1A and BMPR1B by MYC-tagged SMOC2 and mutant SMOC2. Only HA-tagged BMPR1B could be detected in the precipitated immunoprecipitation by MYC-tagged SMOC2 or mutant SMOC2.

According to the above results, we concluded that overexpression of SMOC2 led to too much free SMOC2 competing with BMP2 for binding to BMPR1B. Thus, we assumed that the ability of mutant SMOC2 binding to the matrix proteins declined, and more free mutant SMOC2 could compete with BMP2 for BMPR1B binding (Figure. 7).

**Figure 7.**
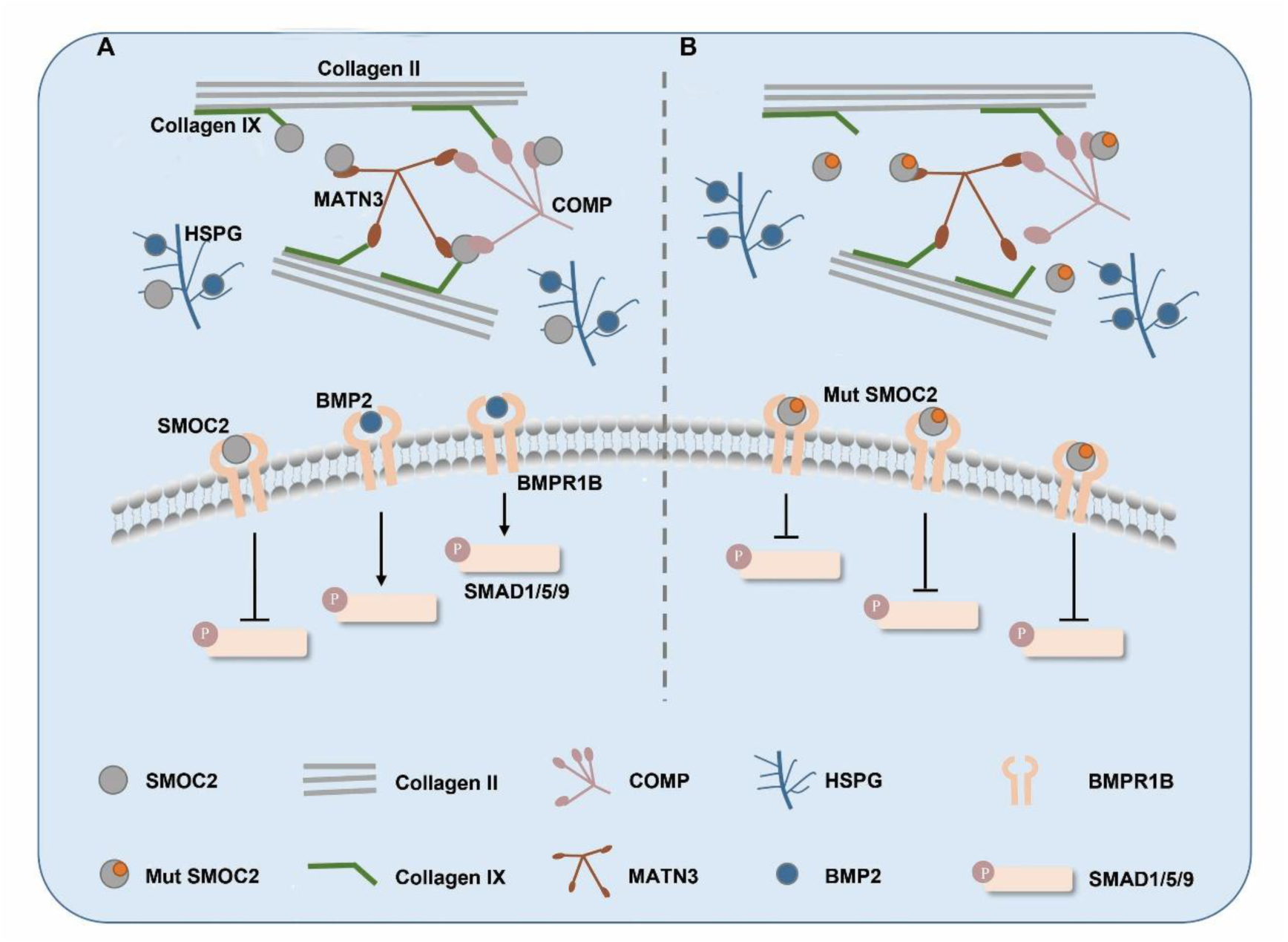
Schematic representation depicting the molecular interactions resulting in negative regulation of BMP-SMAD signaling by wild-type SMOC2 and mutant SMOC2. A: Wild-type SMOC2 could bind to matrix proteins such as COL9A1 and HSPG and competitively bind to BMPR1B with BMP2. B: Mutant SMOC2 bound to matrix proteins weakly, which led to high level of dissociative mutant SMOC2 released into the matrix and more dissociative mutant SMOC2 competing with BMP2 for BMPR1B and phosphorylation of SMAD1/5/9 was blocked.

## Discussion

In this study, we demonstrated a four-generation MED family with complicated phenotypes and identified an unreported pathogenic gene, *SMOC2*, establishing a close relationship of *SMOC2* with MED for the first time. The missense mutation c.1076T>G (p. Leu359Arg) of *SMOC2* was suggested to be the causative variant.

Of note, we detected a meaningful phenotype, the enlarged hypertrophic zone in tibia growth plates in the *Smoc2* knock-in mice model, which had been observed in some knockout mice models, including those deficient in *Matn3*^[15]^, *Smad3*^[16]^, and *Fgf18*^[17]^. The mechanisms of expanded hypertrophic zones are different. The expanded hypertrophic zones in *Matn3*-deficient and *Smad3*-deficient mice were due to accelerated differentiation of chondrocytes^[7][16]^. Both proliferative and hypertrophic zones were expanded in *Fgf18*-deficient mice, which also displayed delayed ossification and decreased expression of osteogenic markers^[17]^. In our study, chondrocyte proliferation did not differ in all mice. Chondrocyte apoptosis was reduced in the hypertrophic zones of tibial growth plates from mutant mice. Furthermore, the increased expression of Ihh in mutant mice may be another reason for the expanded hypertrophic zones. Whether this is related to delayed ossification and matrix resorption needs further investigation.

SMOC2 has been found involved in cell cycle progression by maintaining integrin-linked kinase activity during the G1 phase^[18]^. Both a homozygous mutation (c.84þ1G>T) in the canonical-splice donor site of intron 1 and c.681T>A(p.C227X) nonsense mutations in SMOC2 caused dental development defects ^[19, 20]^. However, no dental defects were observed in our family or the knock-in mouse model. Recent study has shown that overexpressing *SMOC2* in osteoprogenitor cells inhibits osteogenic differentiation and ECM mineralization ^[21]^. *SMOC2* was also considered an antagonist of BMP signaling. Study of zebrafish showed that Smoc2 can inhibit the transcription of BMP target genes ^[22]^. Another study of Xenopus and Drosophila showed that SMOC can activate MAPK signals, thereby inhibiting the BMP signaling downstream of its receptor, and can expand the range of BMP signaling by competing HSPG binding ^[14, 23]^.

SMOC2 and its homologue SMOC1 contain an FS domain, two TY domains separated by one SMOC domain, and an EC domain with two EF-hand calcium-binding motifs family ^[12, 13]^. The EC domain of BM-40 protein families have diverse biological functions such as transducing signals of calcium as a secondary signal, maintaining conformation by binding with calcium, and binding to collagenous proteins^[24]^. Homozygous missense variants in the EC domain of BM-40 abolished the affinity of BM-40 to collagen type I and caused recessive osteogenesis imperfecta type IV^[25]^. Previous study has shown that the EC domain of SMOC2 is fully conserved and is presumably important by binding calcium for the structure of SMOC2^[13, 26]^. The SMOC2 EC domain mediates cell attachment by binding to αvβ6 and αvβ1 integrins on cell surface receptors ^[27]^. In the present report, the Leu359Arg mutant in SMOC2 responsible for MED in our family is strictly conserved in evolution and is located in the first EF-hand motif of the EC domain. When we first focused on the interaction of SMOC2 and collagen IX, we found that the mutation could affect the interaction affinity of SMOC2 and collagen IX. So, the reduced interaction between SMOC2 and collagen IX may alter the biological function of collagen IX or SMOC2 and result in this MED. Certainly, its detailed mechanism needs to be elucidated.

SMOC2 is an important structural anchor in the ECM networks of cartilage, like other members of the BM40/SPARC family, but SMOC2 can also interact with receptors on the cell surface, acting as a regulator of cell–matrix interaction to activate or inactivate some signaling pathways. Previous evidence demonstrated that SMOC2 is involved in the pathways fibroblast growth factor or vascular endothelial growth factor signaling ^[28]^, TGF-β1-SMAD2/3 ^[29, 30]^, WNT/β-catenin ^[31]^, and BMP-SMAD1/5/9 ^[14, 23, 32]^, which play an important role in chondrogenesis. Considering the importance of the BMP-SMAD1/5/9 pathway in endochondral bone formation^[33–36]^, we assessed whether the mutant Smoc2 affected the BMP-SMAD1/5/9 pathway. Our study confirmed that overexpression of wild-type SMOC2 could antagonize BMP signaling, and mutant SMOC2 overexpression still retained this antagonist property intact in vitro, which was due to the mutant site located in the EC domain. In knock-in mice, heterozygous and also homozygous mutant Smoc2 inhibited BMP signaling, which is consistant with wild-type SMOC2 overexpression. So, we conclude that the pathogenic mechanism in this MED family is inhibition of BMP signaling by the mutation (Figure. 7).

The SMOC EC (EC domain only) domain could bind HSPG with similar affinity to BMP2. Low levels of SMOC competitively bound to HSPG and released BMP to bind with BMPR and activate BMP signaling, whereas high amounts of SMOC both bound HSPG and inhibited BMP signaling. We also found that the mutant SMOC2 EC domain reduced or abolished the interaction of SMOC2 with collagen IX and HSPG, which at least led to increased insoluble SMOC2 level in the ECM of cartilage.

In our study, inhibiting the overexpression of SMOC2 to BMP signaling was due to neither increasing degradation of phospho-SMAD 1/5/9 nor MAPK-mediated phosphorylation of the Smad linker region (Figure 5 - figure supplement 1B). Rescue experiments with the BMP receptors ACVR1, caBMPR1A and caBMPR1B further suggests that its inhibition mainly depends on competitive BMP–receptor binding, especially BMPR1B binding, which was not altered by the Leu359Arg mutant in the SMOC2 EC domain. This contrasts with previous study finding that SMOC, mainly SMOC1, acted downstream of the BMPR via MAPK-mediated phosphorylation of the Smad linker region^[14]^.

In conclusion, our evidence supports that *SMOC2* is a new MED-causative gene and illustrates the importance of *SMOC2* in the development of cartilage and long bones. Ultimately, a complete understanding of the molecular genetics and cell-matrix pathophysiology of MED will aid in the diagnosis, prognosis, and treatment of patients and families with MED and will also help in understanding the disease mechanisms of more common conditions such as osteoarthritis.

## Materials and Methods

### Family

We were contacted by a Chinese family that had several adults with short stature and osteoarthritis. All patients in this family received a diagnosis of MED based on clinical and radiography findings. Genomic DNA was extracted from peripheral venous blood by using standard protocols. This study was approved by the medical ethics committees of Shandong University, China and followed the principles of the Declaration of Helsinki. Before the study initiation, written informed consent was obtained from participating individuals.

### Exome sequencing, variant filtration and mutation detection

We performed exome sequencing of DNA from patients II9, III14 and IV9 in this family by using SureSelect Human All Exon Kit (Agilent, Santa Clara, CA) to capture the exome and HiSeq2000 platform (Illumina, San Diego, CA) for sequencing. All variations were filtered by using dbSNP137, the 1000 Genomes Project, and HapMap8 databases. Sanger sequencing was used to confirm the mutation in *SMOC2* in this family. PCR involved using 40 ng genomic DNA and Easy Taq (Transgen Biotech, Beijing). The PCR products were sequenced by Biosune Biotechnology (Shanghai) and compared with the reference sequence in NCBI (https://blast.ncbi.nlm.nih.gov). To predict the detrimental effect caused by the mutation, bioinformatics analysis of the p.Leu359Arg mutation involved using PolyPhen-2, Mutation Taster (http://www.mutationtaster.org/) and SIFT.

### Construction of the targeting vector and generation of chimeric mice

A targeting vector containing SMOC2 exon10, 11 and 12 and the mutation allele flanked by a loxP site and a loxP-neo cassette was constructed, which was introduced into mouse embryonic stem (ES) cells. After removing the cassette, the targeting ES cells were injected into C57BL/ 6 (B6) blastocysts to generate chimeras (F0), which were mated with C57BL/6 mice to generate F1 heterozygous offspring. Generation of the mouse model was performed by Cyagen Biosciences (Guangzhou, China). The genotypes of mice were determined by Sanger sequencing. The sequence containing the mutation was amplified with primers (Table 4). The PCR products were sequenced by Biosune Biotechnology (Shanghai) and compared with the reference sequence in NCBI.

**Table 4.**
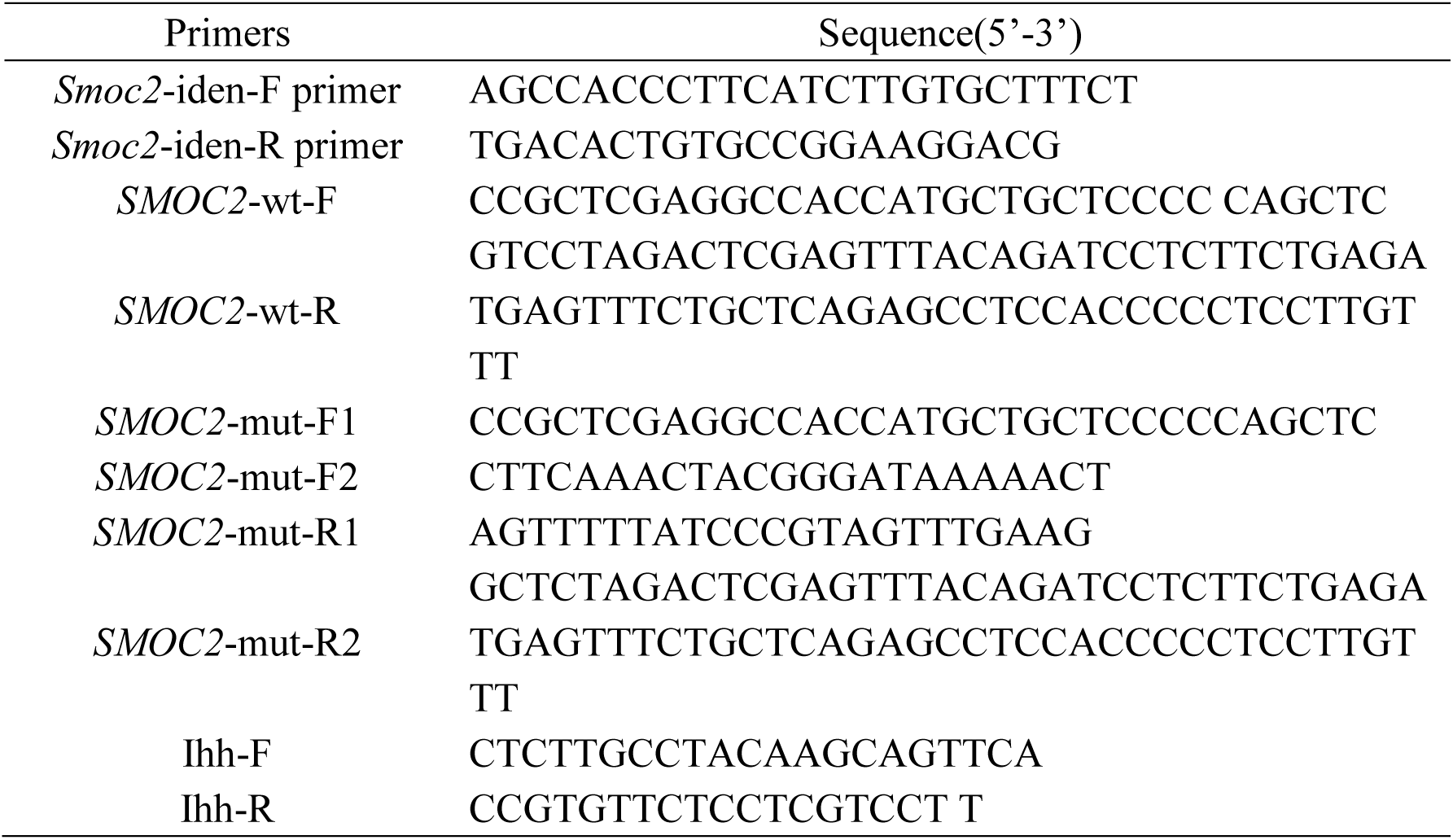
The primers used in this study

### Cell culture

The cells The human embryonic kidney 293T (HEK293T) cell were cultured in DMEM containing 10% FBS (Thermo Fisher Scientific), 100 U/ml penicillin (A603460, Sangon Biotech, Shanghai) and 100 μg/ml streptomycin (A100382-0050, Sangon Biotech) at 37°C with 5% CO2. Primary chondrocytes were taken from the knees of P6 littermates. F-12 nutrient medium was purchased from Gibco (Thermo Fisher Scientific). Cells were cultured in F12 containing 10% FBS at 37°C with 5% CO2.

### Cell transfection and treatment

Recombinant lentiviruses and stably transfected cell line were established as described in previous study^[37]^. Stably transfected cells were seeded in a 100-mm^2^ dish at 80% density. After maintaining in DMEM with 10% FBS overnight, 4–12 μg pCMV3-COMP-HA (HG10173-CY), pCMV3-MATN3-HA (HG11951-UT), or pCMV3-COL9A1-HA (HG12231-CY, all Sino Biological, China); pcDNA3-ACVR1 (80870), pcDNA3-caBMPR1A (80873) or pcDNA3-caBMPR1B (80882); and pcDNA3-ACVR1 (80871), pcDNA3-caBMPR1A (80874) or pcDNA3-caBMPR1B (80883, all Addgene) were transfected into stably transfected cells by using 12 μl PEI (BMS1003-A, Thermo Fisher Scientific). At 48 h after transfection, cells were collected. HEK293 cells stably transfected with or without wild SMOC2 -MYC were incubated in serum-free medium with MG132 (A2585, ApexBio Technology) at increasing doses (10μM, 20μM, 50μM) for 1h and then incubated with BMP at 20 ng/ml for 1h. MYC, p-SMAD1/5/9, p-SMAD1 (Ser206), SMAD1 and GAPDH in whole cell protein lysates were analyzed by western blot.

### Analysis of the skeleton

The body weights of littermates were measured on days 30 and 63. The body length measurements were taken from X-ray radiographs by using X-RAD225 OptiMAX. The lengths of isolated femurs and tibias were measured by using a vernier caliper.

### Alizarin Red and Alcian Blue double-staining

The newborn littermates were sacrificed and the skin, muscle, viscera and soft tissues were removed. After fixation in absolute ethanol and acetone, alcian blue (0.1%, G1027, Servicebio Technology Co. Ltd, Wuhan, China) and alizarin red S (0.2%, G1038, Servicebio Technology) staining of skeleton from newborn littermates was performed.

### Skeleton tissue paraffin section

Hind limbs from littermates at postnatal day 0 (P0), 21 (P21), 63 (P63) and heart, liver, kidney, stomach, duodenum, colon, bladder, cerebrum, cerebellum, gastrocnemius, epiphyseal growth plate of tibia, rib, sternum, vertebra and mandible from *Smoc2^+/+^* mice at P30 were dissected. After fixation overnight in 4% paraformaldehyde and decalcification in 20% EDTA for 1 month, bone samples were embedded in paraffin wax and cut into 4 μm sections.

### Histological analysis

The slides of tissue samples were soaked in 3% H_2_O_2_ (ZLI-9311, ZSGB-BIO, China) for 15 min to quench endogenous peroxidase activity. An amount of 0.1% trypsin (A100458-0050, Diamond) was used to recover antigen and 20% goat serum (ab7481, Abcam) was used for blocking. The 4-μm paraffin-embedded sections of tibia were incubated overnight at 4°C with anti-PCNA (1:20, AB0051, Abways Technology), anti-Ihh (1:100, 13388-I-AP, Cell Signaling Technology) or anti-phospho-Smad1(Ser463/465)/Smad5 (Ser463/465)/Smad9 (Ser465/467) (1:100, 13820S, Cell Signaling Technology) and 1 h with the secondary antibody at room temperature. Then, sections were developed by using 3,3-diaminobenzidine (DAB) (ZLI-9017, ZSGB-BIO). After dehydrating, clearing and mounting, the slices were photographed by microscopy (BX41, OLYMPUS, JPN).

For H&E staining (G1005-100, Servicebio Technology, Wuhan, China), the 4-μm paraffin-embedded sections of tibia were stained in 10% hematoxylin for 5 min and in 1% eosin for 1 min.

For TUNEL analysis, One Step TUNEL Apoptosis Assay Kit (C1090, Beyotime Biotechnology, Shanghai) was used and visualization was by laser-scanning confocal microscopy (BX51, OLYMPUS, Japan). Nuclei were stained with DAPI (ab104139, Abcam) and apoptotic cells were labeled with Cyanine 3 (Cy3).

### Immunofluorescence analysis

For immunofluorescence experiments, monolayers of primary chondrocytes from littermates were soaked in 0.1% TritonX-100 (T0694-100ML, Sangon Biotech) in PBS and blocked with 20% goat serum (ab7481, Abcam), then incubated with antibodies anti-Phospho-Smad1 (Ser463/465)/Smad5 (Ser463/465)/Smad9 (Ser465/467) (1:100, 13820S, Cell Signaling Technology) overnight at 4°C and secondary antibody [Alexa Fluor 594 – conjugated Affinipure Donkey Anti-Rabbit IgG(H+L) antibody (SA0006-8, Proteintech Group)] for 1 h at 37°C in dark. Cell nuclei were stained with 4′,6-diamidino-2-phenylindole (DAPI) (ab104139, Abcam). After sealing with neutral gum, sections were photographed by laser-scanning confocal microscopy (BX51, OLYMPUS, Japan).

### Co-immunoprecipitation (Co-IP)

The proteins from stably transfected cells transiently transfected with *COMP*, *MATN3* or *COL9A1* and *ACVR1*, *BMPR1A* or *BMPR1B* were extracted by using IP lysis buffer with 1% protease inhibitor cocktail (K1007, ApexBio Technology) and ultrasonication. Extracts were mixed with antibody-coated Dynabeads (10004D, Invitrogen, Thermo Fisher Scientific) and incubated at 4°C for 3 h. After 3 washes in washing buffer, immunoprecipitated proteins were eluted in 7.5 μl 2×SDS loading buffer by heating at 99°C for 10 min for western blot analysis.

### Western blot analysis

Total cell lysates or tissue lysates from the P21 mice tissue including heart, lung, liver, muscle, brain, kidney, testis, stomach, duodenum, tibia, lumbar vertebral and sternum were collected in RIPA lysis Buffer (P0013C, Beyotime Biotechnology) with 1% protease inhibitor cocktail (K1007, ApexBio Technology) and 1% phosphatase inhibitor (K1015, ApexBio Technology). Protein concentration was determined according to the instructions of BCA Kit (Boster Biological Technology, Wuhan, China). An amount of 35 or 40 μg whole cell lysates or tissue lysates were resolved by 10% SDS-PAGE and transferred onto PVDF membranes (Merck Millipore, Germany). After incubation in 5% non-fat milk for 1 h at room temperature, the membrane was incubated with the anti-Phospho-Smad1 (Ser463/465)/Smad5 (Ser463/465)/Smad9 (Ser465/467) (1:100, 13820S, Cell Signaling Technology), anti-HA (1:10000, 66006-2-Ig, Proteintech Group), anti-MYC (1:10000, 60003-2-Ig, Proteintech Group), anti-SMAD1 (1:1000, D151630, Sangon Biotech) or anti-GAPDH (1:20000, 60004-1-Ig, Proteintech Group) overnight at 4°C and horseradish peroxidase-conjugated secondary antibody for 1 h at room temperature. Signals were visualized by ECL blotting detection reagents (32209, Thermo) and exposed to X-ray films (SUPER RX-N-C, FUJIFILM).

### Real time PCR (qRT-PCR)

Cartilages from proximal tibias were dissected from wild-type and mutant mice. Then total RNA from cartilages was isolated by using TRIzol (Invitrogen) and cDNA was generated by using random primers. Real time PCR was performed with the SYBR Green Real Time PCR Master Mix (CW0957, CWBIO, CHN) and primers (Supplemental Table S4).

### Solid phase binding assay (SPB)

*SMOC2* gene, SMOC2 FTST domain, SMOC2 EC domain and Mut SMOC2 EC domain containing the His tag were synthesized by Bioss Biotechnology. HSPG (H4777, Sigma-Aldrich) or COL9A1 (TP318130, OriGene Technologies) was coated in the 96-well plates at 40 ng/well. Nonspecific binding sites were blocked with 30% BSA for 2 h at temperature. Wild-type or mutant SMOC2 EC domain (Bioss) was added to the wells at increasing doses (10, 20, 50, 100 and 200 ng/well) and incubated for 2 h at room temperature. Wells were incubated with anti-SMOC2 primary antibody (1:1000, bs-7506R, Bioss) overnight at 4°C and horseradish peroxidase-conjugated secondary antibody for 1 h at room temperature. Then 100 μl TMB (C520026, EL-TMB Chromogenic Reagent kit, Sangon Biotech) was added to every well and plates were kept in the dark until a blue color was obtained. Then, 50 μl stop buffer (C520026, EL-TMB Chromogenic Reagent kit, Sangon Biotech) was added to stop the reaction and OD at 450 nm was measured.

### Transmission electron microscopy (TEM)

TEM was performed by Servicebio Technology. The tibial growth plates from littermates were dissected and fixed in 4% PFA for 4 h. After a wash with 0.1 M sodium cacodylate buffer for 3 times, the samples were fixed with 1% osmium tetroxide for 2 h at room temperature. Samples were dehydrated through ascending ethanol series and ethanol was replaced by acetone. Then samples were embedded with 812 embedding medium and incubated for 48 h at 60°C. Samples were cut at 60-80 nm and stained with uranyl acetate and lead citrate for 15 min and images were obtained by using an HITACHI Transmission Electron Microscope (HT7700).

### Statistical analysis

Unpaired two-tailed Student *t* test was used to analyze the results of the body length and weight of *Smoc2^+/+^, Smoc2^L359R/+^* and *Smoc2^L359R/L359R^* mice at P30 and P63, the length of tibia and femur of *Smoc2^+/+^, Smoc2^L359R/+^* and *Smoc2^L359R/L359R^* mice at P30 and P63, the width of growth plates and the ratio of hypertrophic zone to that of a growth plate of *Smoc2^+/+^, Smoc2^L359R/+^* and *Smoc2^L359R/L359R^* mice at P21 and P63 and immunostaining analysis. P<0.05 was considered statistically significant.

## Acknowledgements

We thank the patients and their families for their participation. This work was supported by the National Natural Science Foundation of China (81671114, 81471602<81741055<81873878) and Natural Science Foundation of Shandong Province (2015GSF118050, ZR2015HZ002, ZR2016HZ01, ZR2012HQ015) and the Key Technology Research and Development Program of Shandong (2016ZDJS07A08, 2018CXGC1211).

---

**Figure 1-figure supplement 1.**
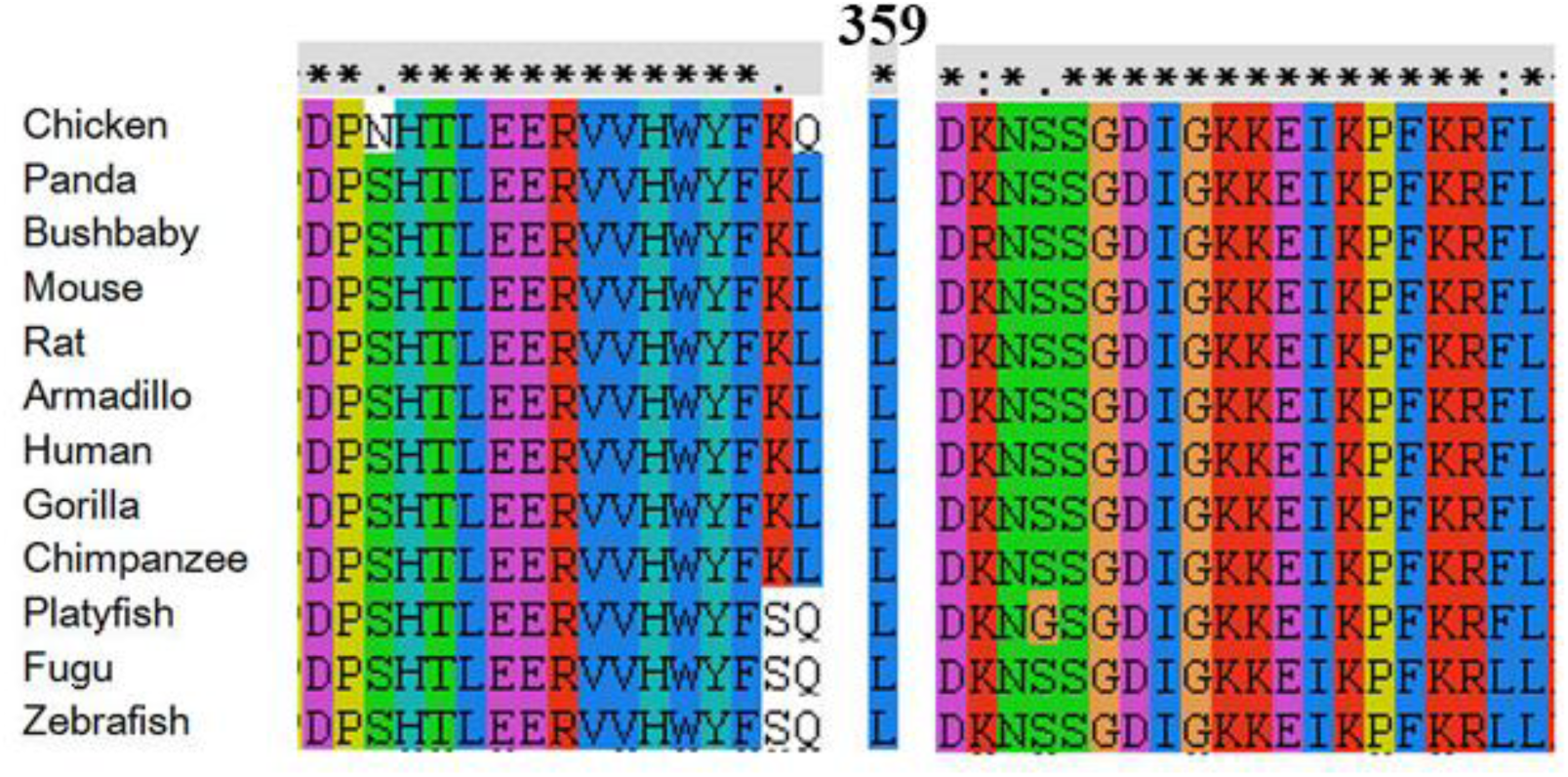
Evolutionary conservation of the leucine at position 359 of SMOC2.

**Figure 2-figure supplement 1.**
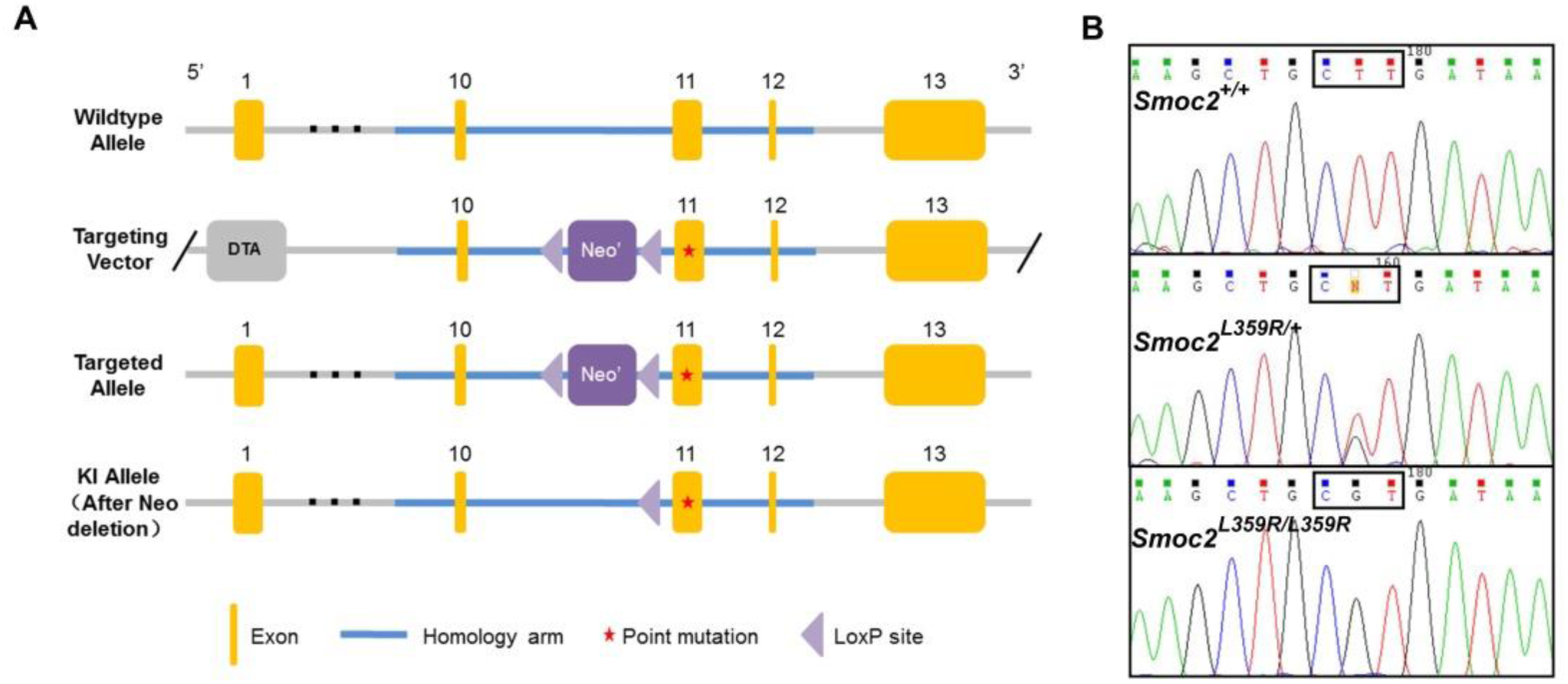
The construction of mutant SMOC2 mouse model. A: The targeting construct with LoxP sequences indicated by arrowheads and the site of the mutation by a red pentagram. B: Sequencing of mutation-positive homologous recombinants confirmed the presence of the mutation (CTT > CGT).

**Figure 2-figure supplement 2.**
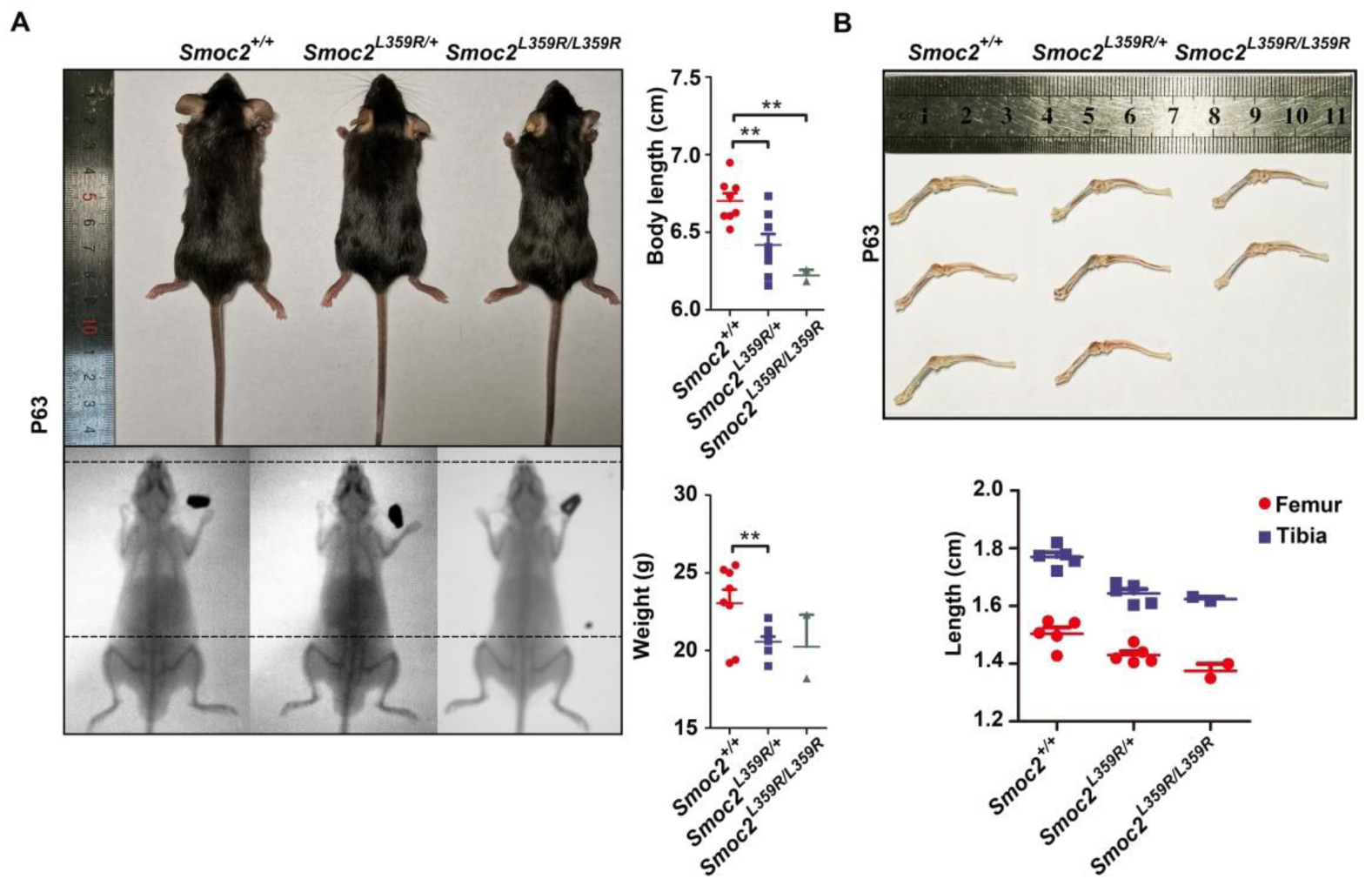
A: Photographs and radiographs of *Smoc2^+/+^, Smoc2^L359R/+^* and *Smoc2^L359R/L359R^* mice at P63. Black dotted lines are aligned at the tip of the nose and the top of the pelvis of the *Smoc2^+/+^* mouse. Body lengths and weights of all mice were measured at P30. (n=8:8:2, ★★P<0.01). B: Lengths of femurs and tibias of all mice were measured at P63.

**Figure 5-figure supplement 1.**
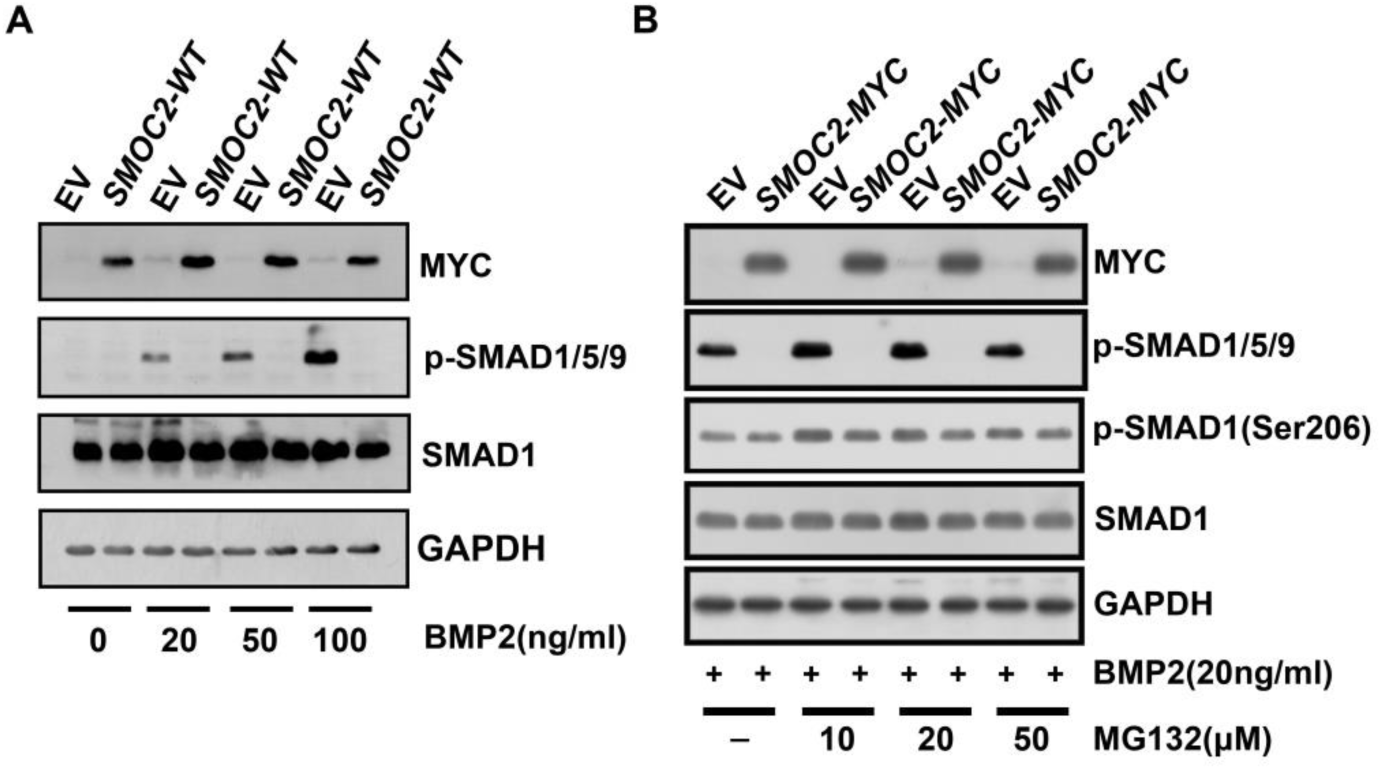
A: HEK293 cells stably transfected with or without wild SMOC2 were incubated in serum-free medium with BMP2 at the indicated doses for 1h. Phosphorylated SMAD1/5/9 (p-SMAD1/5/9), total SMAD1 and GAPDH in whole cell protein lysates were analyzed by western blot. Phosphorylation of SMAD1/5/9 by BMP2 was blocked in cells stably transfected with wild SMOC2. B: HEK293T cells stably transfected with or without wild-type *SMOC2* -MYC were incubated in serum-free medium with MG132 at the indicated doses for 1 h, then incubated with BMP at 20 ng/ml for 1 h. Western blot analysis of MYC, p-SMAD1/5/9, p-SMAD1(Ser206), SMAD1 and GAPDH in whole cell protein lysates.

**Figure 5-figure supplement 2.**
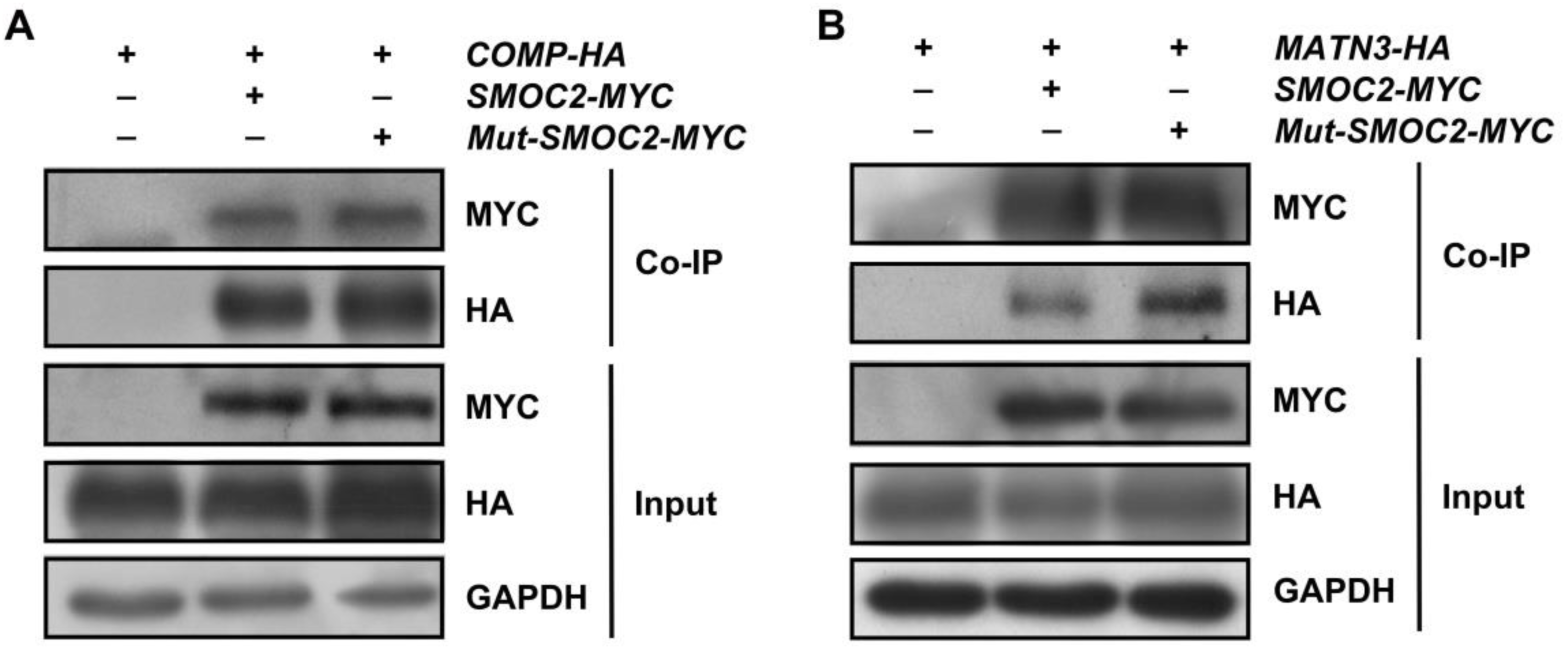
Co-IP showed the interaction of SMOC2 and COMP and MATN3. HA-tagged COMP and HA-tagged MATN3 could be detected in the precipitated immunoprecipitation by MYC-tagged wild-type SMOC2 and MYC-tagged mutant SMOC2

## References

1. Chapman KL, Briggs MD, Mortier GR: Review: clinical variability and genetic heterogeneity in multiple epiphyseal dysplasia. Pediatr Pathol Mol Med 2003, 22(1):53–75.

2. Anthony S, Munk R, Skakun W, Masini M: Multiple epiphyseal dysplasia. J Am Acad Orthop Surg 2015, 23(3):164–172.

3. Briggs MD, Hoffman SM, King LM, Olsen AS, Mohrenweiser H, Leroy JG, Mortier GR, Rimoin DL, Lachman RS, Gaines ES et al: Pseudoachondroplasia and multiple epiphyseal dysplasia due to mutations in the cartilage oligomeric matrix protein gene. Nat Genet 1995, 10(3):330–336.

4. Hecht JT, Nelson LD, Crowder E, Wang Y, Elder FF, Harrison WR, Francomano CA, Prange CK, Lennon GG, Deere M et al: Mutations in exon 17B of cartilage oligomeric matrix protein (COMP) cause pseudoachondroplasia. Nat Genet 1995, 10(3):325–329.

5. Czarny-Ratajczak M, Lohiniva J, Rogala P, Kozlowski K, Perala M, Carter L, Spector TD, Kolodziej L, Seppanen U, Glazar R et al: A mutation in COL9A1 causes multiple epiphyseal dysplasia: further evidence for locus heterogeneity. Am J Hum Genet 2001, 69(5):969–980.

6. Muragaki Y, Mariman EC, van Beersum SE, Perala M, van Mourik JB, Warman ML, Olsen BR, Hamel BC: A mutation in the gene encoding the alpha 2 chain of the fibril-associated collagen IX, COL9A2, causes multiple epiphyseal dysplasia (EDM2). Nat Genet 1996, 12(1):103–105.

7. Paassilta P, Lohiniva J, Annunen S, Bonaventure J, Le Merrer M, Pai L, Ala-Kokko L: COL9A3: A third locus for multiple epiphyseal dysplasia. Am J Hum Genet 1999, 64(4):1036–1044.

8. Chapman KL, Mortier GR, Chapman K, Loughlin J, Grant ME, Briggs MD: Mutations in the region encoding the von Willebrand factor A domain of matrilin-3 are associated with multiple epiphyseal dysplasia. Nat Genet 2001, 28(4):393–396.

9. Dasa V, Eastwood JRB, Podgorski M, Park H, Blackstock C, Antoshchenko T, Rogala P, Bieganski T, Jazwinski SM, Czarny-Ratajczak M: Exome sequencing reveals a novel COL2A1 mutation implicated in multiple epiphyseal dysplasia. Am J Med Genet A 2019, 179(4):534–541.

10. Superti-Furga A, Neumann L, Riebel T, Eich G, Steinmann B, Spranger J, Kunze J: Recessively inherited multiple epiphyseal dysplasia with normal stature, club foot, and double layered patella caused by a DTDST mutation. J Med Genet 1999, 36(8):621–624.

11. Balasubramanian K, Li B, Krakow D, Nevarez L, Ho PJ, Ainsworth JA, Nickerson DA, Bamshad MJ, Immken L, Lachman RS et al: MED resulting from recessively inherited mutations in the gene encoding calcium-activated nucleotidase CANT1. Am J Med Genet A 2017, 173(9):2415–2421.

12. Vannahme C, Smyth N, Miosge N, Gosling S, Frie C, Paulsson M, Maurer P, Hartmann U: Characterization of SMOC-1, a novel modular calcium-binding protein in basement membranes. J Biol Chem 2002, 277(41):37977–37986.

13. Vannahme C, Gosling S, Paulsson M, Maurer P, Hartmann U: Characterization of SMOC-2, a modular extracellular calcium-binding protein. Biochem J 2003, 373(Pt 3):805–814.

14. Thomas JT, Eric Dollins D, Andrykovich KR, Chu T, Stultz BG, Hursh DA, Moos M: SMOC can act as both an antagonist and an expander of BMP signaling. Elife 2017, 6.

15. van der Weyden L, Wei L, Luo J, Yang X, Birk DE, Adams DJ, Bradley A, Chen Q: Functional Knockout of the Matrilin-3 Gene Causes Premature Chondrocyte Maturation to Hypertrophy and Increases Bone Mineral Density and Osteoarthritis. The American journal of pathology 2006, 169(2):515–527.

16. Xiao Yang, Lin Chen, Xiaoling Xu, Cuiling Li, Cuifen Huang, Deng C-X: TGF-beta_Smad3 signals repress chondrocyte hypertrophic differentiation and are required for maintaining articular cartilage. The Journal of Cell Biology 2001, 153(1):35–46.

17. Liu Z, Xu J, Colvin JS, Ornitz DM: Coordination of chondrogenesis and osteogenesis by fibroblast growth factor 18. Genes Dev 2002, 16(7):859–869.

18. Liu P, Lu J, Cardoso WV, Vaziri C: The SPARC-related factor SMOC-2 promotes growth factor-induced cyclin D1 expression and DNA synthesis via integrin-linked kinase. Mol Biol Cell 2008, 19(1):248–261.

19. Bloch-Zupan A, Jamet X, Etard C, Laugel V, Muller J, Geoffroy V, Strauss JP, Pelletier V, Marion V, Poch O et al: Homozygosity mapping and candidate prioritization identify mutations, missed by whole-exome sequencing, in SMOC2, causing major dental developmental defects. Am J Hum Genet 2011, 89(6):773–781.

20. Alfawaz S, Fong F, Plagnol V, Wong FS, Fearne J, Kelsell DP: Recessive oligodontia linked to a homozygous loss-of-function mutation in the SMOC2 gene. Arch Oral Biol 2013, 58(5):462–466.

21. Peeters T, Monteagudo S, Tylzanowski P, Luyten FP, Lories R, Cailotto F: SMOC2 inhibits calcification of osteoprogenitor and endothelial cells. PLoS One 2018, 13(6):e0198104.

22. Mommaerts H, Esguerra CV, Hartmann U, Luyten FP, Tylzanowski P: Smoc2 modulates embryonic myelopoiesis during zebrafish development. Dev Dyn 2014, 243(11):1375–1390.

23. Thomas JT, Canelos P, Luyten FP, Moos M, Jr.: Xenopus SMOC-1 Inhibits bone morphogenetic protein signaling downstream of receptor binding and is essential for postgastrulation development in Xenopus. J Biol Chem 2009, 284(28):18994–19005.

24. Sasaki T, Hohenester E, Gohring W, Timpl R: Crystal structure and mapping by site-directed mutagenesis of the collagen-binding epitope of an activated form of BM-40/SPARC/osteonectin. EMBO J 1998, 17(6):1625–1634.

25. Mendoza-Londono R, Fahiminiya S, Majewski J, Care4Rare Canada C, Tetreault M, Nadaf J, Kannu P, Sochett E, Howard A, Stimec J et al: Recessive osteogenesis imperfecta caused by missense mutations in SPARC. Am J Hum Genet 2015, 96(6):979–985.

26. Novinec M, Kovacic L, Skrlj N, Turk V, Lenarcic B: Recombinant human SMOCs produced by in vitro refolding: calcium-binding properties and interactions with serum proteins. Protein Expr Purif 2008, 62(1):75–82.

27. Maier S, Paulsson M, Hartmann U: The widely expressed extracellular matrix protein SMOC-2 promotes keratinocyte attachment and migration. Exp Cell Res 2008, 314(13):2477–2487.

28. Rocnik EF, Liu P, Sato K, Walsh K, Vaziri C: The novel SPARC family member SMOC-2 potentiates angiogenic growth factor activity. J Biol Chem 2006, 281(32):22855–22864.

29. Luo L, Wang CC, Song XP, Wang HM, Zhou H, Sun Y, Wang XK, Hou S, Pei FY: Suppression of SMOC2 reduces bleomycin (BLM)-induced pulmonary fibrosis by inhibition of TGF-beta1/SMADs pathway. Biomed Pharmacother 2018, 105:841–847.

30. Yuting Y, Lifeng F, Qiwei H: Secreted modular calcium-binding protein 2 promotes high fat diet (HFD)-induced hepatic steatosis through enhancing lipid deposition, fibrosis and inflammation via targeting TGF-beta1. Biochem Biophys Res Commun 2019, 509(1):48–55.

31. Lu H, Ju DD, Yang GD, Zhu LY, Yang XM, Li J, Song WW, Wang JH, Zhang CC, Zhang ZG et al: Targeting cancer stem cell signature gene SMOC-2 Overcomes chemoresistance and inhibits cell proliferation of endometrial carcinoma. EBioMedicine 2019, 40:276–289.

32. DeGroot MS, Shi H, Eastman A, McKillop AN, Liu J: The Caenorhabditis elegans SMOC-1 Protein Acts Cell Nonautonomously To Promote Bone Morphogenetic Protein Signaling. Genetics 2019, 211(2):683–702.

33. Tsumaki N, Nakase T, Miyaji T, Kakiuchi M, Kimura T, Ochi T, Yoshikawa H: Bone morphogenetic protein signals are required for cartilage formation and differently regulate joint development during skeletogenesis. J Bone Miner Res 2002, 17(5):898–906.

34. Retting KN, Song B, Yoon BS, Lyons KM: BMP canonical Smad signaling through Smad1 and Smad5 is required for endochondral bone formation. Development 2009, 136(7):1093–1104.

35. Sanchez-Duffhues G, Hiepen C, Knaus P, Ten Dijke P: Bone morphogenetic protein signaling in bone homeostasis. Bone 2015, 80:43–59.

36. Yoon BS, Ovchinnikov DA, Yoshii I, Mishina Y, Behringer RR, Lyons KM: Bmpr1a and Bmpr1b have overlapping functions and are essential for chondrogenesis in vivo. Proc Natl Acad Sci U S A 2005, 102(14):5062–5067.

37. Ma X, Qiu R, Dang J, Li J, Hu Q, Shan S, Xin Q, Pan W, Bian X, Yuan Q et al: ORMDL3 contributes to the risk of atherosclerosis in Chinese Han population and mediates oxidized low-density lipoprotein-induced autophagy in endothelial cells. Sci Rep 2015, 5:17194.

